# Chemical shift and relaxation regularisation improve the accuracy of ^1^H MR spectroscopy analysis

**DOI:** 10.1101/2025.01.06.631582

**Authors:** Martin Wilson

**Author notes:** **Correspondence** Martin Wilson, Centre for Human Brain Health, University of Birmingham, Edgbaston, Birmingham, B15 2TT, United Kingdom.

## Abstract

**Purpose:** Accurate analysis of metabolite levels from ^1^H MRS data is a significant challenge, typically requiring the estimation of approximately 100 parameters from a single spectrum. Signal overlap, spectral noise and common artefacts further complicate analysis, leading to instability and reports of poor agreement between different analysis approaches. One inconsistently used method to improve analysis stability is known as regularisation, where poorly determined parameters are partially constrained to take a predefined value. In this study we examine how regularisation of frequency and linewidth parameters influences analysis accuracy.

**Methods:** The accuracy of three MRS analysis methods was compaired: 1) ABfit, 2) ABfit-reg and 3) LCModel, where ABfit-reg is a modified version of ABfit incorporating regularisation. Accuracy was assessed on synthetic MRS data generated with random variability in the frequency shift and linewidth parameters applied to each basis signal. Spectra (N=1000) were generated across a range of SNR values (10, 30, 60, 100) to evaluate the impact of variable data quality.

**Results:** Comparison between ABfit and ABfit-reg demonstrates a statistically significant (*p <* 0.0005) improvement in accuracy associated with regularisation for each SNR regime. An approximately 10% reduction in the mean squared metabolite errors were found for ABfit-reg compared to LCModel for SNR *>*10 (*p <* 0.0005). Furthermore, Bland-Altman analysis shows that incorporating regularisation into ABfit enhances its agreement with LCModel.

**Conclusion:** Regularisation is beneficial for MRS fitting and accurate characterisation of the frequency and linewidth variability *in vivo* may yield further improvements.

## INTRODUCTION

^1^H MR spectroscopy (MRS) is a unique tool for measuring brain metabolite levels with numerous applications in both the clinical ^1^ and neuroscience domains ^2,3^. One particularly challenging aspect of MRS is the accurate estimation of metabolite levels amidst confounding factors, such as noise, residual water, outer-volume artefacts and lineshape variability ^4^. Furthermore, these challenges are exacerbated by strong spectral overlap between some metabolite signals. While a number of approaches may be used to extract meaningful information from MRS data ^5^, spectral fitting is currently the most popular and recommended approach for MRS analysis ^6^, and a number of methods have been developed based on this strategy.

The LCModel analysis algorithm has become the most widely used spectral fitting method for MRS research, since its introduction in 1993^7^. The method is based on performing a “least-squares” fit of a “linear-combination” of predefined molecular signals (known as a basis-set) to the experimentally acquired spectral data. The fitting model estimates individual amplitude parameters for each element in the basis set, and following appropriate scaling, these correspond directly to the metabolite levels in the region of interest. In addition to simple scaling amplitudes, a number of non-linear parameters need to be estimated during the fitting process to ensure accurate metabolite level estimates. These include parameters that influence all basis-set signals equally, such as phase and lineshape, and those that are unique to each basis-set element, such as a chemical shift (frequency) and T_2_ relaxation adjustments. These parameters are known to be influenced by experimental conditions, for instance B_0_ inhomogeneity will have a significant influence on all metabolite lineshapes, and physiological changes in temperature and pH are likely to influence individual chemical shifts ^8,9^. Finally, spectral fitting also requires the estimation of the baseline signal, which is characteristically smooth compared to the basis-set signals, and typically arises from incomplete water suppression and scalp lipid contamination. In LCModel, the baseline signal is modelled using smoothing splines ^10^.

More recently, a range of alternative fitting methods have been proposed, which share the same basic principle of the parametric adjustment of a basis set of metabolite signals to match the acquired data in a least-squares sense. These approaches may be broadly characterised based on their baseline modelling strategy, altough we note these methods also differ in a number of other ways. QUEST ^11^ and TARQUIN ^12^ both perform fitting in the time-domain, where the influence of rapidly decaying baseline signals is significantly suppressed by evaluating the fitting residual after a short temporal delay. FSL-MRS ^13^ and OSPREY ^14^ incorporate frequency-domain polynomial and spline baselines respectively, where the baseline smoothness is controlled by adjusting the spectral density of spline knots or highest degree of polynomial function. AQSES ^15^, ABfit ^16^ and ProFit-1D ^17^ all use penalised splines ^18^ for baseline modelling, where baseline smoothness is controlled by a penalty factor parameter, which is automatically determined in ABfit and ProFit-1D. While other MRS fitting approaches have been developed, including those optimised for edited ^19^ and 2D-MRS analysis ^20^, we limit our discussion here to those focused on the analysis of ^1^H MRS data acquired with single-voxel or MRSI localisation schemes — resulting in one spectrum for each spatially encoded location.

MRS fitting algorithm accuracy is typically assessed in two ways: (1) direct comparison with a gold-standard reference method based on experimentally acquired data, or (2) the analysis of synthetic MRS data. Both methods have their relative advantages, with experimental data being better suited to the realistic evaluation of degraded data quality and artefacts on metabolite estimates. However, the “ground-truth” metabolite levels are not available for experimentally acquired data, presenting a significant challenge in judging the relative performance of each method. Knowledge of the genuine metabolite levels is readily available for synthetic MRS, however this comes at the risk of being contrived due to unrealistic synthesis assumptions.

One further strategy for assessing analysis accuracy involves the preparation of a phantom containing metabolites dissolved in solution. Whilst this approach benefits from knowledge of the “ground-truth” concentrations, in practice it is less commonly used than those described above. The preparation of the mixture of around 20 metabolites, required to approximate a typical brain spectrum, can become prohibitively expensive when high accuracy (and therefore chemical purity) is required. Additional complications arise from attempting to match: 1) typical water and metabolite relaxation properties; 2) pH; 3) temperature and 4) macromolecular contributions, to those typically observed *in vivo*.

Larger studies comparing various MRS fitting method have been performed, with Zöllner at al reporting results from a multi-centre dataset comprised of 277 short TE PRESS spectra acquired from healthy participants at 3 Tesla ^21^. Only a weak to moderate agreement was found between approaches, despite relatively high data quality compaired to clinical MRS. A second study, based on synthetic conventional MRS data, also found relatively poor agreement between methods ^22^. Weak agreement between fitting approaches was also found for GABA edited MRS ^23^, a surprising finding considering the relative spectral simplicity of GABA edited spectra compared to conventional short TE MRS.

A typical MRS fitting model, with a basis set of around 25 to 30 signals, requires the estimation of approximately 75 to 100 parameters, in addition to those associated with the baseline model. The stable and accurate estimation of these parameters in the presence of noise, signal overlap and spectral artefacts presents a significant challenge, and overfitting is one likely cause of disagreement between fitting approaches. Restricting parameter estimates to feasible values, for example constraining metabolite amplitude estimates to be greater than or equal to zero, known as “hard-constraints”, is one way to reduce overfitting. Another approach, known as “soft-constraints” or regularisation, encourage parameters to take an expected value. Unlike hard-constraints, soft-constraints may be violated, provided a clear deviation is supported by the data. For example, the default behaviour of LCModel enforces a soft-constraint on the level of GABA to take the value of 0.04 times the sum of total-NAA, total-creatine and 3×total-choline. Whilst soft-constraints of this type improve fitting stability for signals with comparable intensities to the noise level, they have also been shown to introduce unwanted bias, and are generally not advised ^22^. Furthermore, bias introduced by these soft-constraints is likely exacerbated in pathology — where metabolite levels are more variable, e.g. low or absent total-NAA in brain tumour spectra.

Regularisation may also be applied to the parameters used to model the small frequency and linewidth differences between the basis-set signals and acquired data. Unlike amplitude regularisation, as described above, the relationship between fitting accuracy and regularisation of these non-linear parameters has not yet been reported. In this study, the ABfit fitting method is compared to a new version, incorporating regularisation of the frequency and relaxation parameters (ABfit-reg), and LCModel using synthetic MRS data. Regularisation is shown to significantly improve accuracy for the expected variations in frequency and relaxation, and ABfit-reg is shown to provide either comparable, or improved accuracy relative to LCModel.

## 2 METHOD

### 2.1 Regularisation implementation

Support for the regularisation of the individual basis signal frequency and relaxation parameters is added to ABfit algorithm ^16^, available as part of the spant ^24^ MRS analysis package. To differentiate between the closely related approaches we will use the terms “ABfit” to denote the method as described previously ^16^, and “ABfit-reg” to denote the updated version with regularisation. All aspects of the ABfit and ABfit-reg methods are identical except for those explicitly stated below.

During the final phase (step 4) of the ABfit algorithm ^16^ small individual frequency shift (**f**_*i*_) and Lorentzian line-broadening adjustments, due to T_2_ relaxation, (**d**_*i*_) are applied in the time-domain to each signal in the basis-set **M**^TD^:

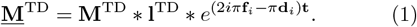

A “global” lineshape adjustment term (**l**^TD^) is also applied equally to all signals in the basis-set to model inhomogeneity in the static magnetic field. The global lineshape model is Gaussian, modified with an asymmetry parameter (*a*_*g*_), as described by Stancik and Brauns ^25^. Transforming the modified basis to the frequency domain 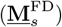 and combining with a basis of spline functions (**B**), to model smooth baseline features, leads to the modelled spectrum **ŷ**:

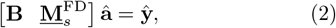

where **â** represents the combined vector of amplitudes for each signal in the basis and spline function. The objective function is defined as the least-squares difference between the modelled spectrum (**ŷ**) and the acquired data 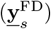, and non-linear fitting parameters, e.g. frequency and linewidth, are adjusted to minimise this function using the Levenberg-Marquardt algorithm ^26^. In contrast to ABfit, terms to regularise the frequency shifts 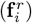, Lorentzian line-broadening 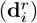 and global lineshape asymmetry 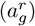 parameter are appended to the objective function in ABfit-reg as follows:

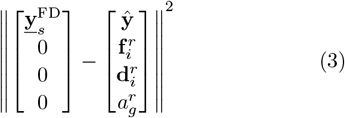

where the double vertical bars represent the *l*^2^ norm. Equation 4 defines the non-regularised objective function used by ABfit for comparison:

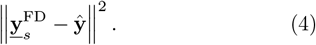

Each of the penalty terms added to equation 3 follow the basic form of scaling the regularised fitting parameters by the standard deviation of the noise (*σ*), estimated from a signal free region of the acquired spectrum, and scalar values determining the regularisation strength for frequency shifts (*f*_reg_), Lorentzian line-broadening (*d*_reg_) and global lineshape asymmetry (*a*_reg_):

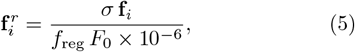

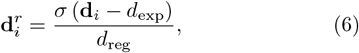

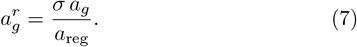

This specific form of regularisation was chosen to be compatible with the approach taken by LCModel, where the regularisation strength is scaled to have comparable magnitude to the spectral noise. Frequency shift regularisation is scaled by *F*_0_ *×* 10^*−*6^ to convert to ppm units, where *F*_0_ is the transmitter frequency in Hz. The expected Lorentzian line-broadening in Hz parameter (*d*_exp_) is introduced to penalise deviations from this common value, whereas values deviating from zero are penalised for the frequency shift and global lineshape asymmetry parameters.

### 2.2 Synthetic MRS

Synthetic MRS data were generated using the spant analysis package ^24^ to establish the influence of regularisation on MRS fitting accuracy. Spectra were generated from a basis set comprised of 29 signals, and corresponding concentrations, listed in Table 1. Metabolite concentrations were based on approximate ranges for healthy brain tissue as listed in de Graaf ^27^. Metabolite simulations were based on chemical shift and J-coupling values from Govindaraju et al ^28^, whereas lipid and macromolecular resonances were based on parameters in Table 1 from Wilson et al ^12^. An additional inverted singlet resonance (-CrCH2) at 3.913 ppm was included in the basis set, as it is often used to model the effect of water suppression on the downfield creatine and phosophocreatine CH2 resonances. The basis set was simulated for the semi-LASER pulse sequence ^29^ at a field strength of 3 Tesla (*F*_0_ = 127.8 *×* 10^6^ Hz) and an echo-time of 28 ms (TE1=8 ms, TE2=11 ms, TE3=9 ms) with ideal RF pulses. Each basis signal was simulated over 1024 complex points sampled at a temporal frequency of 2000 Hz.

**TABLE 1.** Basis set signals and synthetic MRS data concentrations.

| Name (abbreviation) | Conc. (mM) |
| --- | --- |
| Alanine (Ala) | 0.2 |
| Aspartate (Asp) | 1.5 |
| Creatine (Cr) | 5.0 |
| Gamma-aminobutyric acid (GABA) | 1.0 |
| Glucose (Glc) | 1.0 |
| Glutamate (Glu) | 10.0 |
| Glutamine (Gln) | 3.0 |
| Glutathione (GSH) | 2.0 |
| Glycerophosphocholine (GPC) | 1.2 |
| Glycine (Gly) | 1.0 |
| Lactate (Lac) | 0.5 |
| Myo-inositol (Ins) | 5.0 |
| N-acetyl aspartate (NAA) | 10.0 |
| N-acetyl aspartyl glutamate (NAAG) | 2.0 |
| Phosphocholine (PCho) | 0.5 |
| Phosphocreatine (PCr) | 4.5 |
| Phosphoethanolamine (PEth) | 0.8 |
| Scyllo-inositol (sIns) | 0.4 |
| Taurine (Tau) | 1.5 |
| -CrCH2 | 0.0 |
| Lip09 | 0.0 |
| Lip13a | 0.0 |
| Lip13b | 0.0 |
| Lip20 | 0.0 |
| MM09 | 4.0 |
| MM12 | 4.0 |
| MM14 | 4.0 |
| MM17 | 4.0 |
| MM20 | 4.0 |

Prior to summation of the basis signals, normally distributed random Lorentzian linebroadening and frequency shifts were applied. Assumptions about the expected variability in linebroadening and frequency shifts were derived from the LCModel manual ^30^ to ensure a fair comparison between methods. The standard deviation of frequency shifts was 0.004 ppm, with a mean of zero, equivalent to the LCModel parameter DESDSH. The expected Lorentzian broadening is derived from the DEEXT2 parameter as follows:

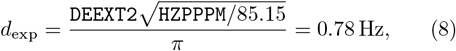

where HZPPPM = *F*_0_ *×* 10^*−*6^ and DEEXT2 = 2.0. Similarly, the standard deviation of the Lorentzian broadening is determined as:

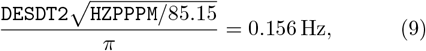

where DESDT2 = 0.4. Following the application of random linebroadening and frequency shifts, basis signals were scaled according to the concentrations listed in Table 1, summed, and dampened with 4 Hz Gaussian linebroadening to model inhomogeneity in the static magnetic field typically observed for good quality MRS data acquired at 3 Tesla. Normally distributed random complex noise was added to generate sets of spectra with different SNRs.

1000 spectra with differing noise samples, frequency shifts (**f**_*i*_) and linebroadening parameters (**d**_*i*_) were generated across 4 spectral SNR (10, 30, 60 and 100) regimes. SNR was calculated as the maximum spectral value divided by the standard deviation of the noise, measured from the real component of the spectra. A second set of data were also generated in the same way across the 4 SNR regimes, except frequency shifts and linebroadening parameters followed a uniform (rather than normal) statistical distribution. Minimum and maximum limits for the uniformly distributed parameters were set to 95% confidence intervals of the standard deviations used for the normally distributed dataset: *±*0.004 *×* 1.96 ppm for the frequency shifts and 0.78 *±*0.156 *×* 1.96 Hz for linebroadening.

Water reference data was also generated, consisting of a singlet resonance at 4.65 ppm with a Gaussian linewidth of 4 Hz and an amplitude (25116) corresponding to the default assumptions for LCModel water-scaling ^30^ (`WCONC = 35880, ATTH2O=0.7`).

The 8 synthetic MRS datasets, comprised of 1000 spectra each, basis-set and water reference data, are available from Zenodo https://doi.org/10.5281/zenodo.14165738 in NIfTI MRS format ^31^. Code to generate the synthetic data, and reproduce all results presented in the following sections, is available from GitHub https://github.com/martin3141/abfit_reg_paper.

### 2.3 Fitting method details and comparison

Synthetic MRS data was analysed with three different methods: 1) ABfit as previously described ^16^; 2) ABfit-reg as described above and 3) LCModel ^7^. A comparison between ABfit and ABfit-reg was performed to isolate the specific influence of regularisation, since all other aspects are identical. LCModel was also included to investigate if regularisation improved the agreement between fitting methods.

The default LCModel method was applied with the NRATIO fitting parameter set to zero, disabling soft-constraints on metabolite ratios, as recommended by Marjańska et al ^22^ to avoid the potential for bias. The following regularisation parameters were used with ABfit-reg to allow a direct comparison with the assumptions made by LCModel: *f*_reg_ = 0.004 ppm, *d*_exp_ = 0.78 Hz, *d*_reg_ = 0.156 Hz. The lineshape regularisation parameter was set as: *a*_reg_ = 0.1, approximately equal to the standard deviation of the 95% confidence interval of *±*0.25 used in ABfit. Upper and lower constraints for **f**_*i*_ and *a*_*g*_ were not enforced in ABfit-reg and non-negativity was the only constraint for **d**_*i*_. Finally, constraints on the the global frequency offset parameter (*f*_0_), in the step 4 of the algorithm, were increased from a negligibly small value in ABfit to 0.05 ppm in ABfit-reg.

A fourth exploratory analysis was performed where the ABfit-reg method was used as described above, but with an additional hard constraint applied to the individual frequency shifts (10^*−*6^ *>* **f**_*i*_ *> −*10^*−*6^), effectively eliminating these parameters from the fitting model. This, “fixed frequency”, fitting approach will be referred to as ABfit-reg-ff.

Fitting accuracy was evaluated as the sum of squared differences from the known concentrations across the 19 metabolite signals listed in Table 1. Agreement between ABfit-reg and LCModel was assessed using Bland-Altman plots ^32^ and all statistical tests were performed in the R software environment ^33^.

## 3 RESULTS

Example plots of fitting results for three example spectra (SNR=10, 30, 100) for ABfit, ABfit-reg and LCModel are shown in Figure 1. The consistent absence of spectral features above the noise level in the fit residual and flat baselines suggest a similarly high level of analysis quality between the approaches.

**FIGURE 1.**
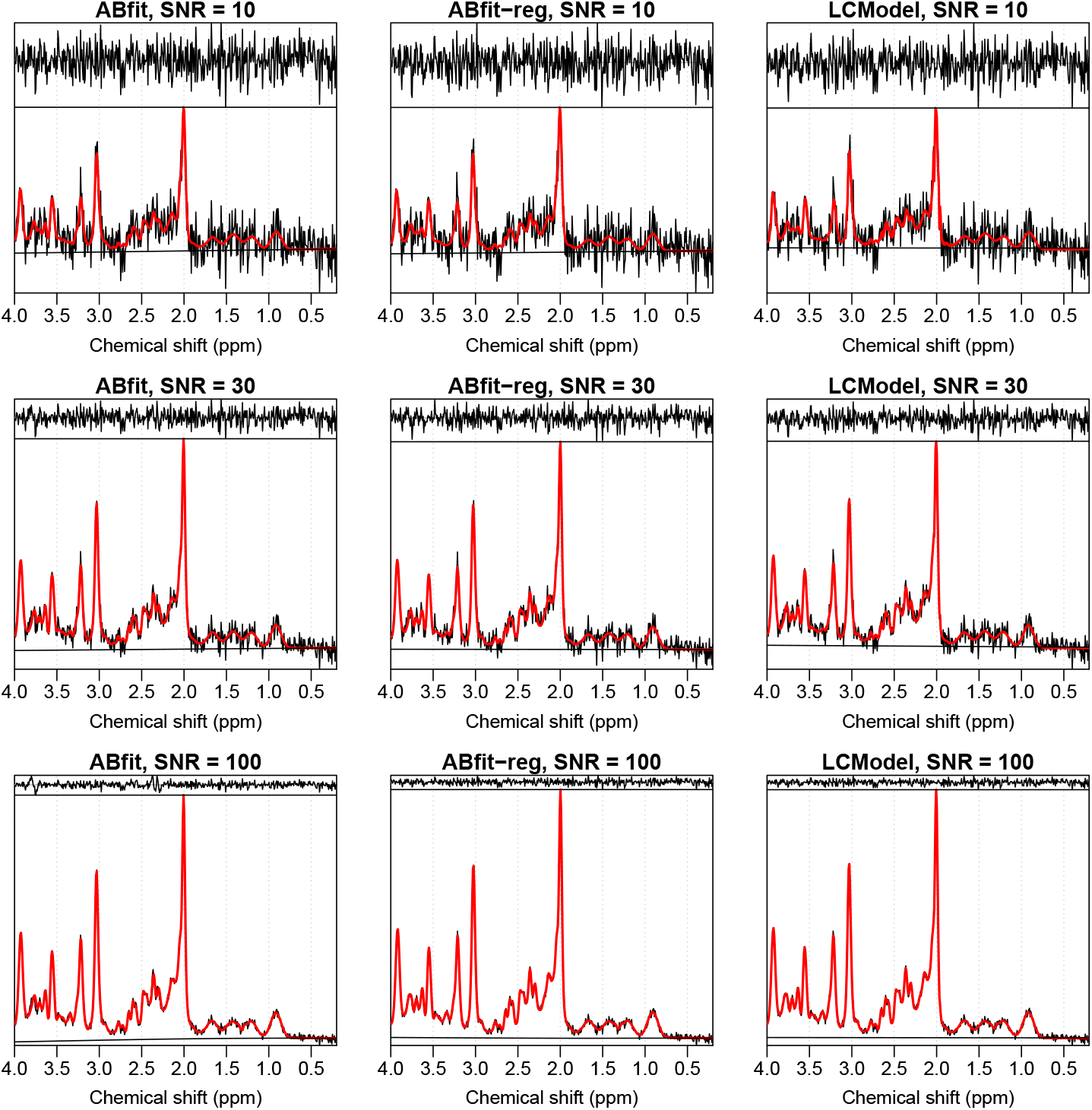
Fitting result plots of three example spectra (SNR=10, 30, 100) analysed with ABfit, ABfit-reg and LCModel. Acquired spectral data is shown in black with the fit shown in red. The black traces above and below the acquired spectrum represent the fitting residual and estimated baseline respectively.

Figure 2 shows a clear difference between the accuracy of ABfit and ABfit-reg, despite the similar appearance in fitting results (Figure 1). The addition of regularisation results in a statistically significant improvement in accuracy of ABfit across all of the four SNR regimes investigated. The relative reduction in the mean squared metabolite error between ABfit and ABfit-reg was 22%, 31%, 45% and 60% for SNRs of 10, 30, 60 and 100 respectively. The reducing relative error associated with increasing SNR suggests that errors associated with local-minima may be dominant at higher SNR, and that regularisation guides the optimisation procedure to improved solutions.

**FIGURE 2.**
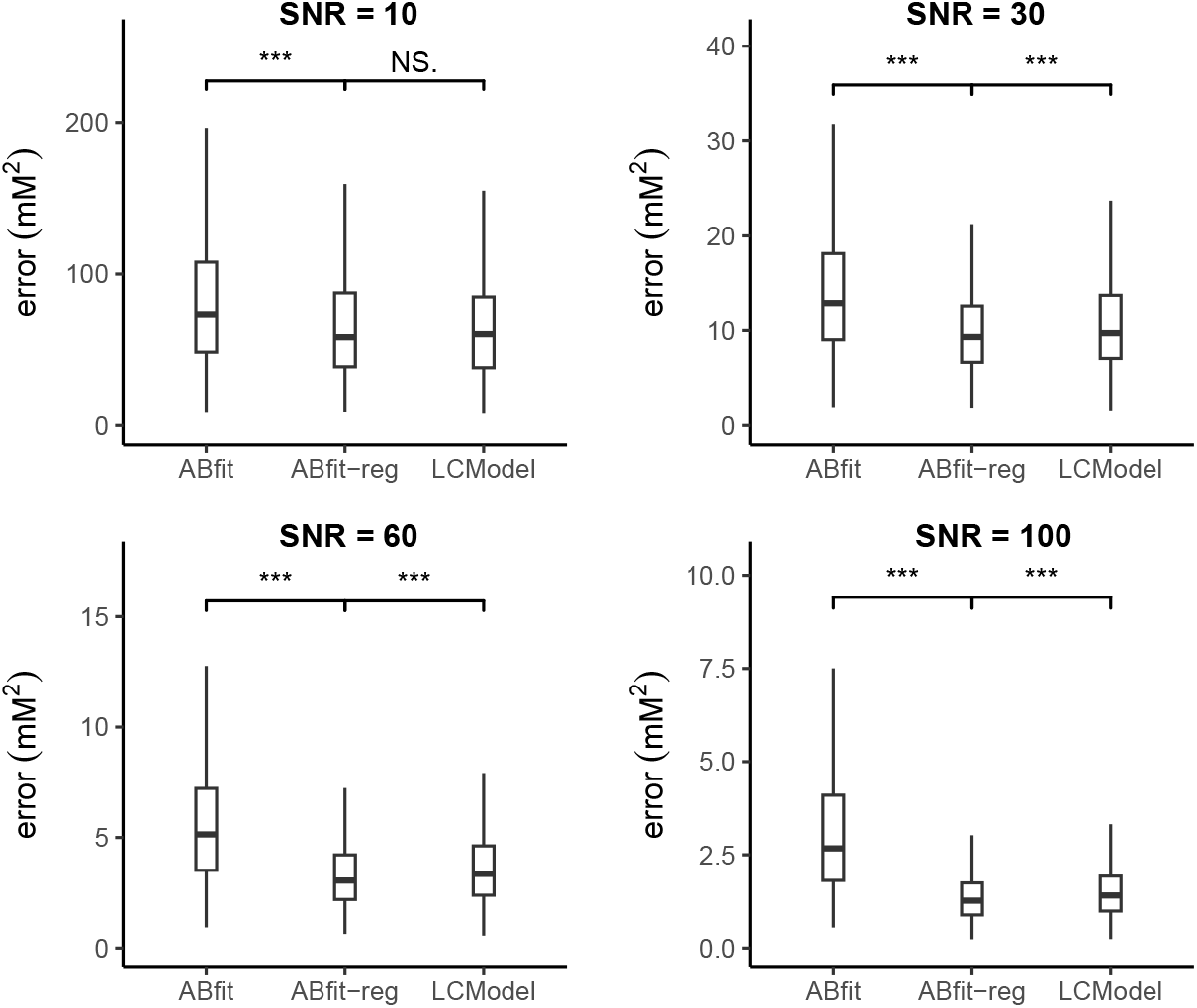
Box and whisker plots comparing metabolite estimate errors (sum of squared differences) for ABfit, ABfit-reg and LCModel from 1000 spectral fits per spectral SNR regime. Normally distributed random frequency shift and linebroadening parameters were applied to individual basis signals. Outliers are not plotted to aid visual comparison betweenthe median values. Statistical significance labels represent a t-test between the two fitting methods: NS. not significant; * *p <*0.05; ** *p <* 0.005; *** *p <* 0.0005.

ABfit-reg is shown to have a statistically significant improvement in accuracy over LCModel for SNRs of 30, 60 and 100, and that the methods have effectively the same accuracy for an SNR of 10 (Figure 2). The relative reduction in the mean squared metabolite error between LCModel and ABfit-reg was 8%, 10% and 11% for SNRs of 30, 60 and 100 respectively.

Influence of the statistical distribution of random shifts and damping factors is very minor (Figure 2 vs S1), suggesting the advantages of regularisation are not heavily dependant on the assumption of normality.

Figure S2 shows that individual frequency shift regularisation has comparable accuracy to simply fixing these shifts to zero (**f**_*i*_ = 0) at SNR = 10. This is likely due to the errors associated with spectral noise being comparable, or greater, than the errors associated with inaccurate individual frequency estimation. For SNRs greater than 10, the errors associated with poor frequency alignment become dominant, with ABfit-reg demonstating improved accuracy over ABfit-reg-ff (*p <* 0.0005).

Bland-Altman analysis was performed to establish if regularisation improved agreement between fitting methods. Figure 3 shows the agreement between LCModel and ABfit / ABfit-reg for GABA, Glu and tNAA (NAA + NAAG), chosen due to their differing contributions to typical healthy brain MRS. Strong reductions in both bias and variability are observed for ABfit-reg vs LCModel compaired to ABfit vs LCModel, demonstrating that regularisation is a significant factor to consider when comparing fitting results from experimentally acquired data - where the true accuracy is unknown.

**FIGURE 3.**
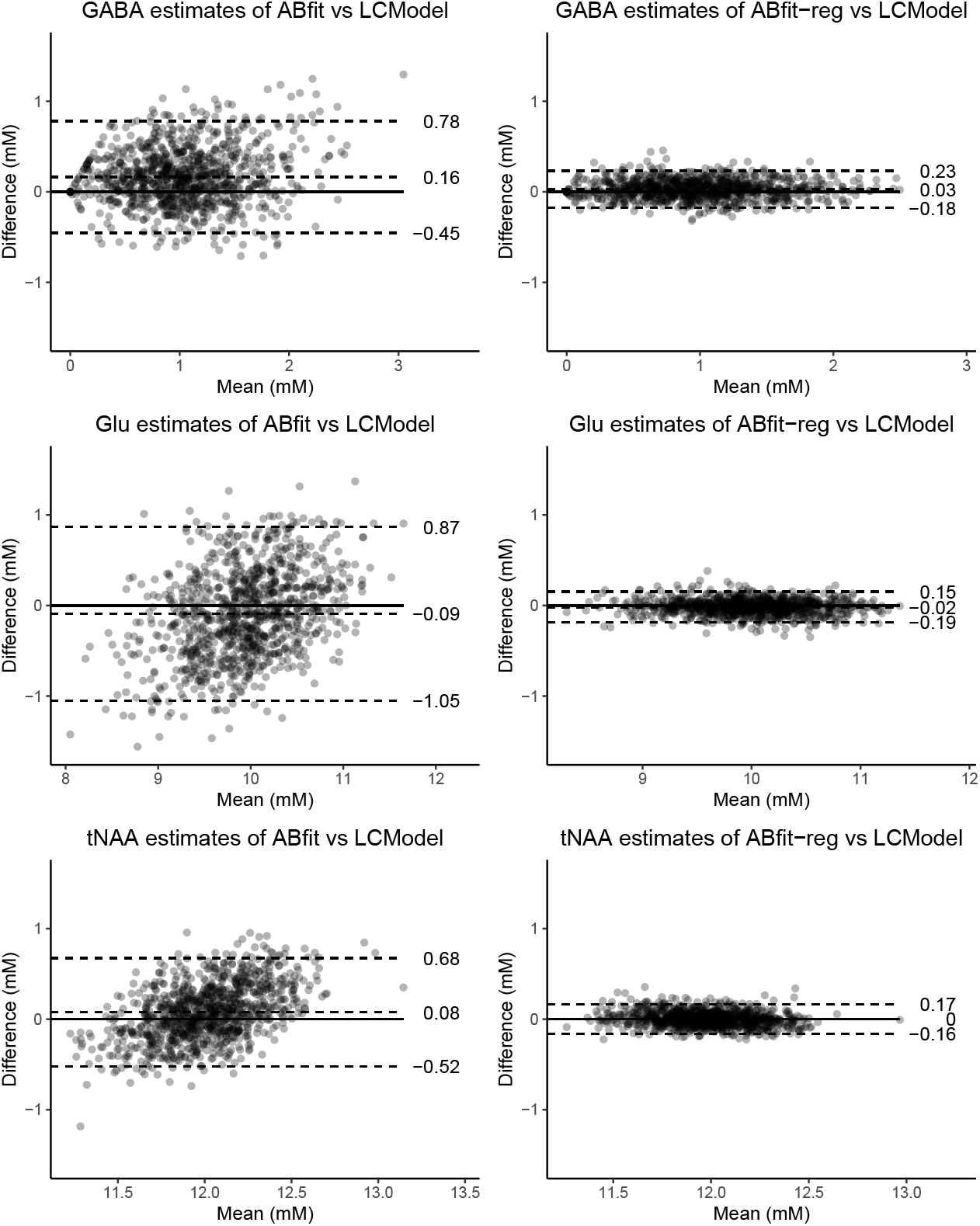
Bland-Altman plots comparing agreement between regularised (ABfit-reg, LCModel) and non-regularised (ABfit) fitting methods. The central horizontal dashed line represents the mean difference, with lower and upper dashed lines representing the 95% confidence intervals of mean difference.

## 4 DISCUSSION

Regularisation of the non-linear fitting parameters for small adjustments to the frequency and linewidth of each basis signal is an integral part of the widely-used LCModel fitting method. However, to the best of the author’s knowledge, a direct evaluation of the benefits of this approach have not been reported. In this study, we adapt the ABfit method to incorporate regularisation (ABfit-reg) and demonstrate an improvement in fitting accuracy across a range of spectral SNRs. Furthermore, we show that incorporating regularisation into ABfit enhances its agreement with LCModel, offering a partial explanation for the generally poor agreement between different fitting approaches ^21,22,23^.

We also note a small, but statistically significant, improvement in accuracy of ABfit-reg compaired to LCModel for spectral SNRs greater than 10. One potential explanation for this contrast in performance may be due to the different approaches to estimate baseline smoothness, however baseline distortions were intentionally not included in the synthetic data model to reduce bias from this aspect. Another possibility is the global lineshape model in LCModel has a greater level of flexibility compaired to ABfit-reg. Since the simulated data model assumes a simple Gaussian lineshape, we may expect ABfit-reg to have improved accuracy due to less degrees of freedom (two lineshape parameters), whereas LCModel may be expected to perform better for more heavily distorted lineshapes due to its “model-free” approach.

In this simulation study we make the same assumptions about random frequency and linewidth changes as LCModel to ensure a fair comparison. However the LCModel manual ^30^ states these assumptions are based on expected differences between *in vivo* and *in vitro* conditions. Historically, metabolite basis sets for MRS analysis were experimentally acquired from spherical phantoms containing solutions of individual metabolites *in vitro*. However, since the development of accurate numerical spectral simulation ^34,35^ and published chemical shift and J-coupling metabolite values ^28^, the use of simulated basis has become the standard approach. Experimentally acquired basis sets are typically acquired at room temperature, whereas simulation parameters are derived from experiments acquired at the more useful physiological temperature of (37°C). Since metabolite chemical shifts have a known dependence on temperature ^8^, it is possible that the range of frequency deviations assumed by LCModel are overestimated for simulated basis sets. Furthermore, chemical shifts are also known to depend on pH ^9^, therefore we might expect lower variability for normally functioning brain tissue compaired to ischemic or cancerous tissue. Further work could be undertaken to measure the true variability of metabolite signals *in vivo*, for healthy and diseased tissue, to optimise regularisation strength — potentially resulting in more accurate metabolite measures.

In this study, we focused on the most commonly used *in vivo* MRS spectral fitting model, where each metabolite corresponds to one signal, modified by a single frequency shift parameter and single linebroadening parameter. Whilst commonly used, this approach is a simplification of the potential variability in metabolite resonances, where each set of chemically equivalent spins (in the same molecule) has the potential for independent frequency and linebroadening terms. These, more complex, models have been developed primarily for *in vitro* analysis of diseased tissue ^36^, where larger variations in pH necessitate greater model freedom, and higher SNR reduces the risk of overfitting. The use of such models *in vivo* is more controversial, since the evidence for significant “proton-group” independence is less established and lower SNR and spectral resolution increase the potential for overfitting. Suggested future work involves estimating the magnitude of these “proton-group” changes from high quality *in vivo* data, and the use of regularisation to reduce fitting instability when applying these more complex models to poorer quality data.

In conclusion, we have demonstrated the benefits of regularisation for MRS analysis, and highlight the potential of characterising frequency and linewidth variability *in vivo* for further improvements in accuracy.

## Data availability statement

Synthetic MRS datasets are available from Zenodo https://doi.org/10.5281/zenodo.14165737. Code to generate the synthetic data and reproduce all figures and results is available from GitHub https://github.com/martin3141/abfit_reg_paper.

