## supporting information for "Chemical shift and relaxation regularisation improve the accuracy of ^1^H MR spectroscopy analysis"

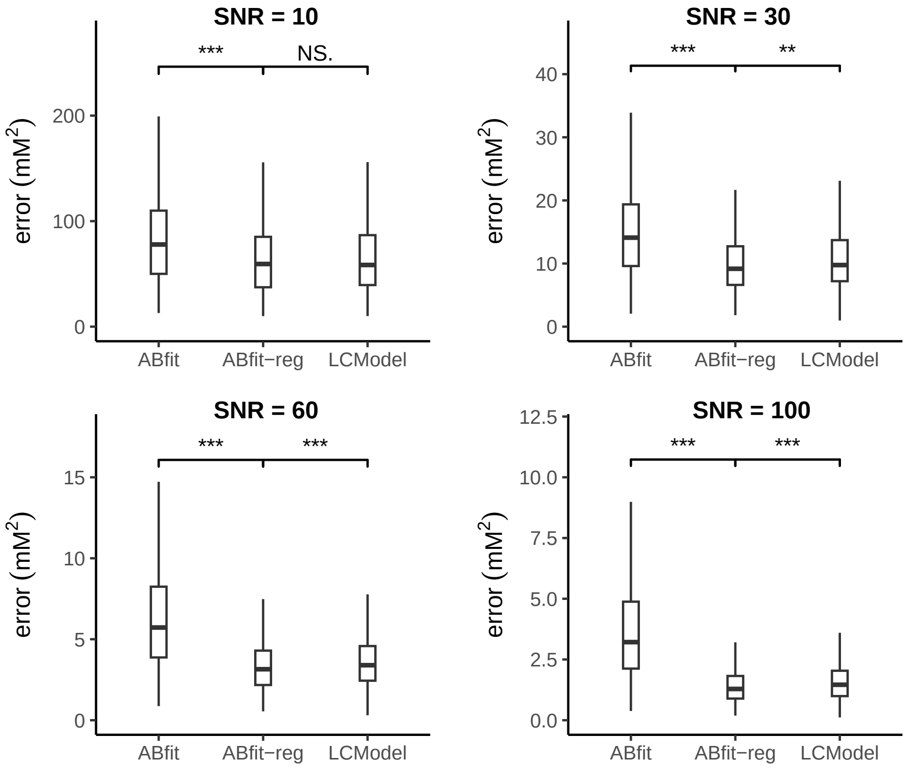


**FIGURE S1** Box and whisker plots comparing metabolite estimate errors for ABfit, ABfit-reg and LCModel from 1000 spectral fits per spectral SNR regime. Uniformly distributed random frequency shift and linebroadening parameters were applied to individual basis signals. Outliers are not plotted to aid visual comparison between the median values. Statistical significance labels represent a t-test between the two fitting methods: NS. not significant; * $p < 0.05$; ** $p < 0.005$; *** $p < 0.0005$.

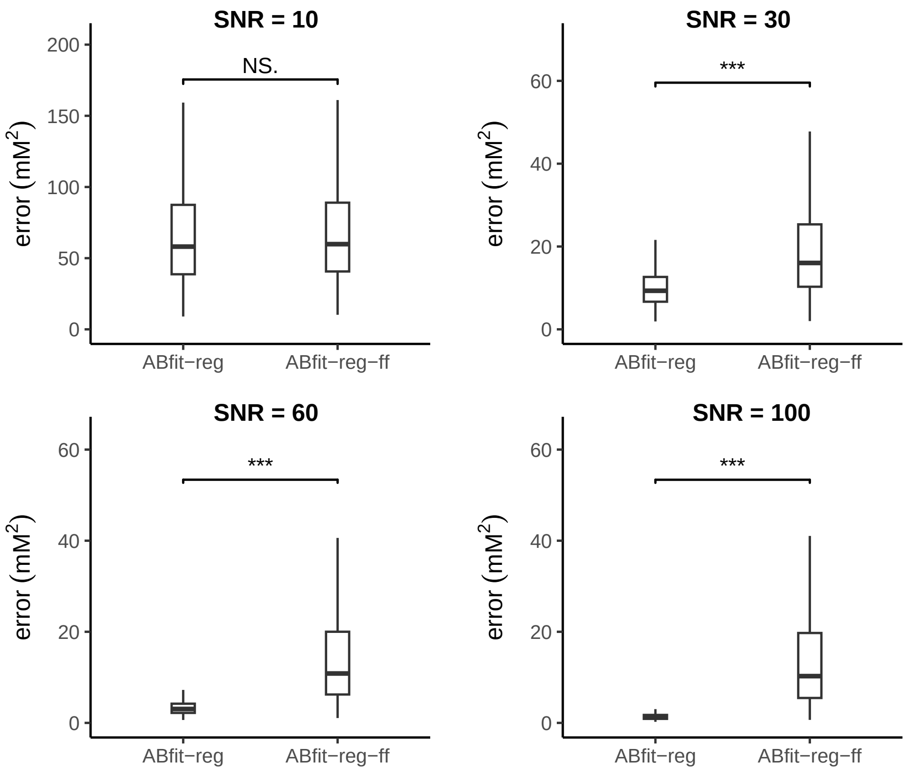


**FIGURE S2** Box and whisker plots comparing metabolite estimate errors for ABfit-reg and ABfit-reg with fixed ($\mathbf{f}_i = 0$) individual frequency parameters (ABfit-reg-ff). Plots are derived from 1000 spectral fits per spectral SNR regime. Normally distributed random frequency shift and linebroadening parameters were applied to individual basis signals. Outliers are not plotted to aid visual comparison between the median values. Statistical significance labels represent a t-test between the two fitting methods: NS. not significant; * $p < 0.05$; ** $p < 0.005$; *** $p < 0.0005$.
